# SCAR FORMATION AND DECREASED CARDIAC FUNCTION FOLLOWING ISCHEMIA/REPERFUSION INJURY IN 1-MONTH-OLD SWINE

**DOI:** 10.1101/773218

**Authors:** Emma J Agnew, Nivedhitha Velayutham, Gabriela Matos Ortiz, Christina M Alfieri, Victoria Moore, Kyle W Riggs, R. Scott Baker, Aaron M Gibson, Sithara Raju Ponny, Tarek Alsaied, Farhan Zafar, Katherine E Yutzey

**Affiliations:** Molecular Cardiovascular Biology, Cincinnati Children’s Hospital Medical Center, 3333 Burnet Ave, Cincinnati, OH, 45229; Cincinnati Children’s Hospital Heart Institute, Department of Pediatrics, University of Cincinnati College of Medicine Cincinnati, OH, 45229; Division of Pediatric Cardiothoracic Surgery, The Heart Institute, Cincinnati Children’s Hospital Medical Center, University of Cincinnati, Cincinnati, OH, 45229; Division of Human Genetics, Cincinnati Children’s Hospital Medical Center, 3333 Burnet Avenue, Cincinnati, OH, 45229

**Author notes:** Corresponding Author: Katherine E. Yutzey, PhD, Division of Molecular Cardiovascular Biology, Cincinnati Children’s Medical Center ML7020, 240 Albert Sabin Way, Cincinnati, OH 45229. Contribution to study: Designed research: E.J.A, R.S.B, K.W.R, T.A, F.Z, K.E.Y. Execution: E.J.A, N.V, G.M.O, C.M.A, V.M, R.S.B, K.W.R, A.G, S.R.P, F.Z. Interpretation: E.J.A, N.V, G.M.O, C.M.A, V.M, K.W.R, S.R.P, T.A, F.Z, K.E.Y. Prepared figures: E.J.A, S.R.P, K.E.Y. Drafted manuscript: E.J.A, K.E.Y. Edited and approved final manuscript: all authors.

**Keywords:** Ischemia/reperfusion injury, cardiac function, cell cycling, cardiomyocyte, pig

## Abstract

Studies in mice show a brief neonatal period of cardiac regeneration with minimal scar formation. Less is known about reparative mechanisms in large mammals. A transient cardiac injury approach (ischemia/reperfusion, IR) was used in weaned postnatal day (P)30 pigs, to assess regenerative repair in young large mammals. Female and male P30 pigs were subjected to cardiac ischemia (1 hour) by occlusion of the left anterior descending artery followed by reperfusion, or to sham operation. Following IR, myocardial damage occurred, with cardiac ejection fraction significantly decreased 2 hours post-ischemia. No improvement or worsening of cardiac function to the 4 week study end-point was observed. Histology demonstrated cardiomyocyte (CM) cell cycling at 2-months-of-age in multinucleated CMs in both sham-operated and IR pigs. Regional scar formation and inflammation in the epicardial region proximal to injury were observed 4 weeks post-IR. Sex differences were found, suggestive of females creating a greater fibrotic response with worse cardiac function, highlighting the importance of representing both sexes in cardiac injury studies. Together, our results describe an effective novel cardiac injury model in P30 swine, at a time when CMs are still cycling. Pigs subjected to IR show myocardial damage with a prolonged decrease in cardiac function, formation of a small, regional scar with increased inflammation. These data demonstrate that P30 pigs do not regenerate myocardium, even in the presence of CM mitotic activity, but form a scar after transient IR injury.

**NEW & NOTEWORTHY:** Here, we report for the first time ischemia/reperfusion (IR) cardiac injury in 1-month-old (P30) pigs. This model of IR injury highlights lack of cardiac regeneration, even in the presence of cardiomyocyte (CM) cell cycling in young swine. An effective injury approach is described for use in large mammals to investigate cardiac function, CM cell cycling, extracellular matrix (ECM) remodeling, and gene expression changes, while highlighting the importance of studying both sexes.

## INTRODUCTION

The ability of the heart to regenerate has been demonstrated in zebrafish, amphibians and neonatal mammals (49). In mice and pigs, a brief neonatal window of cardiac regenerative capacity after injury has been linked to the proliferation of cardiomyocytes (CM)(34, 51, 54). However, whether or not the human heart can regenerate, alongside the timing and mechanisms involved in loss of cardiac regenerative capacity in mammals, are unknown. Thus, there is an unprecedented need to further research cardiac development and repair following injury in the large mammalian setting. Here, we examine the ability of the young swine heart to repair following ischemia/reperfusion (IR) injury.

In mammals, the prenatal heart typically grows by hyperplasia, with mitosis including cytokinesis of mononucleated CMs (9). Following birth, the heart grows primarily by CM hypertrophic growth and, in mice, CMs undergo cell cycle arrest within 7 days (21, 34, 42). This coincides with CMs becoming binucleated, with a consequent increase in cell ploidy, which is then maintained throughout adulthood (21, 46, 47). It is known that mice can regenerate their hearts with pre-existing CMs being the source of regenerating cardiac muscle following injury in the first week after birth (34). However, this intrinsic capacity for cardiac self-repair is lost by postnatal day (P)7 coincident with CM binucleation, cell cycle arrest, and the transition to hypertrophic growth (12, 34, 35, 43). There is incomplete information available as to how the transient regenerative capacity of the neonatal mouse heart specifically relates to CM cell cycle exit, transition to hypertrophic growth, sarcomeric maturation, and fibrotic scar formation in response to injury (28, 34, 35).

In large mammals, less is known of the timing of CM proliferation, multinucleation, hypertrophic growth, and regenerative capacity. Studies in large mammals are required for clinical translation to humans, with thorough analysis of proliferation, nucleation and ploidy across multiple postnatal stages. Recent studies from our group (50) found that the postnatal pig heart shows robust CM mitotic activity in the first 2-months after birth, peaking at P15, resulting in longitudinal growth of CMs by multinucleation. Sarcomeric maturation, multinucleation and hypertrophic CM growth continues to 2-6 months after birth in swine, with CM nuclei predominantly diploid from P0-6 months of age. The presence of significant numbers of mononucleated diploid CMs in pig hearts in the month after birth, alongside active cardiac cell cycling, suggests that there may thus be an extended period for postnatal cardiac regenerative repair via CM proliferative mechanisms after injury in swine (32).

Recent reports have demonstrated a limited regenerative period of 2-3 days after birth in neonatal pigs subjected to permanent ligation of the LAD, which leads to a severe drop in cardiac function (51, 54). Studies in adult swine show no evidence of cardiac regeneration after ischemic injury without intervention, and previous reports in pigs are primarily focused on developing therapeutic targets to aid in cardiac repair (1, 10, 18, 25). In addition, there are inconsistencies in the use of female and/or male animals in previously reported swine cardiac injury studies. Here we report the cardiac injury response in female and male pigs after transient ischemic injury at P30 when CM mitotic activity, multinucleation, and myocardial remodeling are ongoing in postnatal swine.

## MATERIALS AND METHODS

All animal experiments conform to NIH guidelines and were performed with protocols approved by the Institutional Animal Care and Use Committee at the Cincinnati Children’s Research Foundation. White Yorkshire-Landrace farm pigs were purchased from Isler Genetics (Prospect, OH). All animals (n=19 total, with n=16 surviving the full protocol) were aged 1 month (P30) at time of surgery, following 2-3 days of acclimatization upon arrival at CCHMC animal facility. The weight range of pigs at time of surgery was 6-10kg.

### Ischemia/reperfusion (IR) cardiac injury

P30 pigs were randomly assigned to sham (n=8) or IR (n=11) cardiac surgery groups. Left ventricular myocardial samples were also collected from unoperated 2-month old pigs (n=4), to be used as control samples. Of the eleven P30 pigs subjected to IR injury, eight survived the day of the surgery and no subsequent morbidity or mortality was observed. Equal numbers of males (n=4) and females (n=4) were evaluated in each group. On the day of surgery, anesthesia was induced with intramuscular injection of ketamine (Zoetis, Kalamazoo, MI) and xylazine (Akorn Animal Health, Lake Forest, IL). Each pig was intubated with an endotracheal tube of 4-5 French diameter, with anesthesia being sustained by 1-2% isoflurane (Akorn Animal Health) with 100% oxygen. Continuous pulse oximetry measured blood oxygenation throughout the procedure. Three-lead electrocardiogram (ECG) was used to monitor for arrhythmias. A rectal probe was inserted to measure body temperature.

Prior to incision, amiodarone was given intramuscularly (1mg/kg, APP Pharmaceuticals, Schaumburg, IL). IV fluids were maintained at 30 cc/hr with 5% dextrose in Lactated Ringer’s solution. Left lateral thoracotomy was performed to approach the 4^th^ intercostal space and the 3^rd^ or 4^th^ rib was removed after examining the heart’s position. After opening the pericardium, a 4-0 Prolene suture (Ethicon, NJ) was placed around the LAD artery just distal to the 2^nd^ diagonal branch. All IR pigs were administered heparin (1,000 units, Sagent Pharmaceuticals, Schaumburg, IL) and a rubber snare was used to occlude the LAD. ST segment changes were observed on the ECG and the myocardium was visually examined for ischemic changes distal to the spot of occlusion. LAD occlusion lasted 1 hour, after which the snare was released and reperfusion was monitored for 30 minutes. For the sham operations, the LAD suture was not snared down. The chest was left open for 1.5 hours in total, comparable to IR operations. In all pigs, an air knot was left to mark the site of injury for accurate tissue collection at euthanasia. Epinephrine (1 mg) and lidocaine (2ml/mg)(Henery Schein, Dublin, OH), cardiac defibrillation with 10J+ energy, and cardiac massage were given appropriately and as needed in all animals who developed significant arrhythmias bradycardia during the procedure. For chest closure, the ribs were re-opposed with 2-0 Vicryl stitches and tissue closed in layers with Vicryl and Monocryl sutures (Ethicon, NJ and Medline, IL), followed by 1-2 mg/kg of Bupivicaine (Hospira, Lake Forest, IL) was administered around the incision site.

Peripheral blood samples were collected before surgery and 2 hours following IR (or sham equivalent) to measure physiological parameters, including blood chemistry and circulating cTnI, indicative of myocardial damage, via the iSTAT blood analyzer (Abbott Laboratories Diagnostics, IL). Antibiotics were administered by intramuscular injection of Combi-Pen-48 (30,000U/Kg, Bimeda, Oakbrook Terrance, IL) on the day of and 2 days after surgery. Pain management medication was induced with intramuscular injection twice a day for two days with Buprenorphine Hydrochloride (0.01mg/lg, PAR Pharmaceuticals, Chestnut Ridge, NY).

Two pigs in the IR group did not survive the procedure, following bradycardia at 20 and 40 minutes respectively post-ischemia. One IR pig showed poor recovery post-surgery and was euthanized 7-hours post-ischemia. All animals that survived the day of surgery survived the full study (n=16). One month after surgery, pigs were euthanized, via injection of Fatal-Plus (1cc per 4.5kg, Vortech Phamaceuticals, Dearborn, MI), for tissue collection and processing.

### Echocardiography

Transthoracic echocardiography was performed prior to surgery, 2 hours post-cardiac suture placement, 1 week, 2 weeks, and 4 weeks post-operatively using the General Electric Vivid 7 system (Boston, MA) with Digiview software, cardiac package (Tech Tools, Rowlett, TX). Briefly, pigs were anaesthetized as described above (intramuscular injection of ketamine, followed by 1-2% isoflurane with 100% oxygen after intubation), placed in left lateral decubitus position with electrocardiogram electrodes attached to monitor heart rate and a rectal probe inserted to observe body temperature. The sonographer was blinded to the experimental groups during all imaging and data analysis. Images were acquired, with a 5 MHz probe, in 2D mode short and long axis, with M-mode images taken in short axis. Tissue Doppler Imaging measurements of the septal and lateral wall of the left ventricle (LV) were made. 2D parasternal long axis four chamber images were analyzed for LV diastolic and systolic dimensions (cm) and volumes (ml), stroke volume (ml), fractional shortening (%) and ejection fraction (%). M-mode images taken at the mid-papillary muscle end were utilized for analysis of left ventricular septum diastolic and systolic thickness (cm) and LV posterior wall diastolic and systolic thickness (cm). Heart rate in beats per minute (bpm) was obtained from M-mode images. Digiview analysis software was used for calculating all above parameters.

### Histology and tissue staining

For tissue processing, 3 zones (approx. 1 cm cubes) of P30 pig myocardium 4 weeks post-surgery were collected. The scar zone was defined as the region directly inferior to the occlusion site where loss of blood flow and myocardial ischemic damage would be expected to occur. Upon processing, it was noted this area contained both scar tissue and adjacent myocardium in IR pigs. In sham pigs, the same size piece directly at the occlusion site was taken and named ‘scar’ for regional comparisons. The border zone was defined as the area of the LV adjacent to the area classified as ‘scar’. The remote zone was taken from the posterior side of the middle LV. Myocardium was fixed in 4% Paraformaldehyde (PFA) (Electron Microscopy Sciences, Hatfield, PA) in Phosphate-buffered saline (PBS) (Fisher Scientific, Hampton, NH), then transferred to 70% ethanol after 48 hours. Tissues were processed, embedded in paraffin wax and cut using a microtome to 5μm sections of the myocardium, encompassing both the endo- and epicardial regions.

#### Masson’s Trichrome

Masson’s Trichrome 2000 Stain Kit (American Master Tech Scientific, McKinney, TX) was used to identify collagen and muscle in all myocardial sections, according to manufacturer’s procedures. Brightfield images were obtained with an Olympus BX51 microscope and Nikon DS-Ri1 camera (both Tokyo, Japan). Deposition of collagen was quantified on an automated analysis program created on NIS elements software (Tokyo, Japan).

#### Immunohistochemistry

Sections were dewaxed with xylene (Fisher Scientific, Hampton, NH) and rehydrated in decreasing concentrations of ethanol (Decon Labs Inc, PA). Antigen retrieval was carried out in citrate buffer (pH 6; Vector, Burlingame, CA) using a pressure cooker. Slides were blocked with 6% goat serum, followed by staining with Anti-phospho-Histone H3 (Ser10) Antibody (pHH3, 1:100; 06-570), in combination with the cardiomyocyte marker Anti-Troponin I Antibody (cTnI, 1:1000; MAB1691) (both Millipore Sigma, Burlington, MA). Fluorophore-conjugated secondary antibodies (1:100; Alexa Fluor ab175473, ab150081, Abcam, Cambridge, UK) were used to detect immunostaining, with the addition of Wheat Germ Agglutinin (WGA), Alexa Fluor™ 647 Conjugate (1:250; Thermo Fisher Scientific, Waltham, MA) and DAPI (5mg/ml, D1306; Thermo Fisher Scientific). Immunofluorescence was imaged using a Nikon Eclipse Ti Fluorescence microscope with NIS elements software (Tokyo, Japan). The percent pHH3-positive cardiomyocytes, normalized to cardiomyocyte nuclear number, was calculated using an automated analysis program created on NIS elements software. Fiji (ImageJ) image analysis software was used to manually draw the WGA-defined circumference of cardiomyocytes with a central nucleus to calculate cross sectional area (μm^2^).

#### Collagen Hybridizing Peptide detection

Slides were dewaxed and rehydrated before addition of Streptavidin reagent (Component A) for 30 minutes at 37°C, followed by Component B after washing (E21390, Thermo Fisher Scientific). Slides were blocked with 5% goat serum and Collagen Hybridizing Peptide (CHP) solution (15μM, BIO300, 3Helix, Salt Lake City, UT) prepared by heating to 80°C before placing in ice for 15 seconds prior to addition to tissue sections for overnight incubation at 4°C. Streptavidin, Alexa Fluor™ 568 conjugate secondary antibody (1:300, Thermo Fisher Scientific) was added and tissues imaged as above. CHP was calculated as area (μm^2^) per field.

### RNA analysis

Myocardial tissue samples from all three zones isolated in parallel with histology samples described above were snap frozen in liquid nitrogen and stored at −80°C. RNA was extracted using the NucleoSpin RNA kit (740955, Macherey-Nagel, Duren, Germany). The manufacturer’s protocol was followed, with elution in 40μl RNase-free H_2_0. RNA was treated with DNase I treatment (EN0255, Thermo Fisher Scientific, Waltham, MA), according to manufacturer’s protocols. cDNA was synthesized, using the SuperScript III First-Strand Synthesis SuperMix kit (Thermo Fisher Scientific), then subjected to Real-Time quantitative PCR (RT-qPCR) in duplicate using the Step One Plus system (Applied Biosystems, Foster City, CA) with gene-specific primer sets (Supplementary Table 1) and Applied Biosystems Power SYBR Green PCR Master Mix (Thermo Fisher Scientific). Porcine primers were designed on Primer-BLAST (NCBI National Center for Biotechnology Information). Primers were tested for specificity using Sanger DNA sequencing (performed by CCHMC DNA Core) of pooled cDNA sample PCR products, following purification with use of the QIAquick PCR Purification Kit (Qiagen, Hilden, Germany). Relative quantification was provided by StepOne software v2.3 using the quantitation-relative standard curve method with 40 cycles followed by melt curve production. mRNA levels were normalized to the mean *18S* rRNA levels, used as internal standards, and expressed relative to unoperated 2-month-old pig myocardial samples.

#### RNA-sequencing

RNA sequencing (RNA-seq) was performed using the Illumina HiSeq 2500 system in collaboration with the CCHMC DNA Core. Total RNA from pig myocardial tissue was subjected to the TruSeq mRNA LS Illumina protocol with a GeneAmp 9700 Applied Biosystems thermocycler. All sample/library quality control analysis was carried out on the AATI Fragment Analyzer (Agilent) and quantified using a Qubit fluorimeter (Thermo Fisher Scientific). Adapter dimers in the libraries were removed from the pool using a 1.5% gel and cleaned using the QiaQuick Gel Extraction protocol (Qiagen). Sequencing was performed by TrueSeq polyA stranded selection with 75 bp paired sequencing to extract 20 million reads. The samples analyzed included eight samples per sham and eight samples per IR injury, with both scar and border zone samples of the myocardium (n=16 total, n=4 per zone per surgery). Both female and male pigs were used (n=2 per zone per surgery group).

#### Gene expression data analysis

RNA-seq data analysis was carried out by CCHMC Division of Biomedical Informatics. The FASTQ files were obtained from the DNA Sequencing and Genotyping Core facility at CCHMC. Quality control steps were performed to determine overall quality of the reads from the FASTQ files. Upon passing basic quality matrices, the reads were trimmed to remove adapters and low-quality reads using Trimmomatic (2).

The trimmed reads were mapped to the *Sus Scrofa* (swine) reference genome. Hisat2 was used for reference alignment. In the next step, transcript/gene abundance was determined using “Kallisto” (3). A transcriptome index in Kallisto using Ensemble cDNA sequences for the pig was created. This index was then used to quantify transcript abundance in raw counts and transcripts per million (TPM). The quantified sample matrix was used for determining differential gene expression between experimental groups or to profile expression of a specific transcript across various conditions. The R package RUVSeq (40) was used to perform differential gene expression analysis between groups, with raw counts obtained from Kallisto used as input. Significant differentially expressed genes were obtained using a fold change cutoff of 2 and adjusted p-value/p-value cutoff of ≤0.05. The data files were uploaded to the GEO database (accession: GSE137293).

#### Gene Ontology and Pathway Analysis

Downstream functional annotation of genes of interest and significantly dysregulated genes in the experiment were determined using gene ontology (GO) (cellular components, molecular function and biological process) and pathway analysis. A detailed functional annotation and pathway analysis was performed using ToppFun tool from the ToppGene Suite (4). An adjusted p-value cutoff of ≤0.05 was used to select functional annotations and pathways.

### Statistical analysis

All numerical data are presented as means ± SD, with p<0.05 deemed significant. GraphPad Prism 8 software was used for statistical analysis, with Mann-Whitney, Kruskal-Wallis with Dunn’s post-hoc analysis, two-way ANOVA with Tukey post-hoc analysis, or Mixed-Effect Model (for repeated measures) with the Geisser-Greenhouse correction with Bonferroni post-hoc tests, performed as appropriate (as indicated in each Figure legend). All data were subject to Shapiro-Wilk normality testing prior to further analysis.

## RESULTS

### Cardiac ischemia/reperfusion (IR) injury, via temporary occlusion of the LAD, produces myocardial tissue damage in P30 pigs

Since CM cell cycling rates are similar at P0 and P30 in young swine (50), cardiac repair following temporary injury was assessed in P30 pigs. Blood collection and echocardiography were performed at baseline and subsequent time points over the course of the 4 week study (Figure 1A). On the day of the surgery, 3 out of 11 IR pigs did not survive (Figure 1B), due to cardiac arrest during the procedure or poor recovery following the cardiac injury. An additional 2 pigs required resuscitation interventions during surgery (Table 1). Following day 1 of the study, all IR and sham pigs then survived to the end of the protocol.

**Figure 1:**
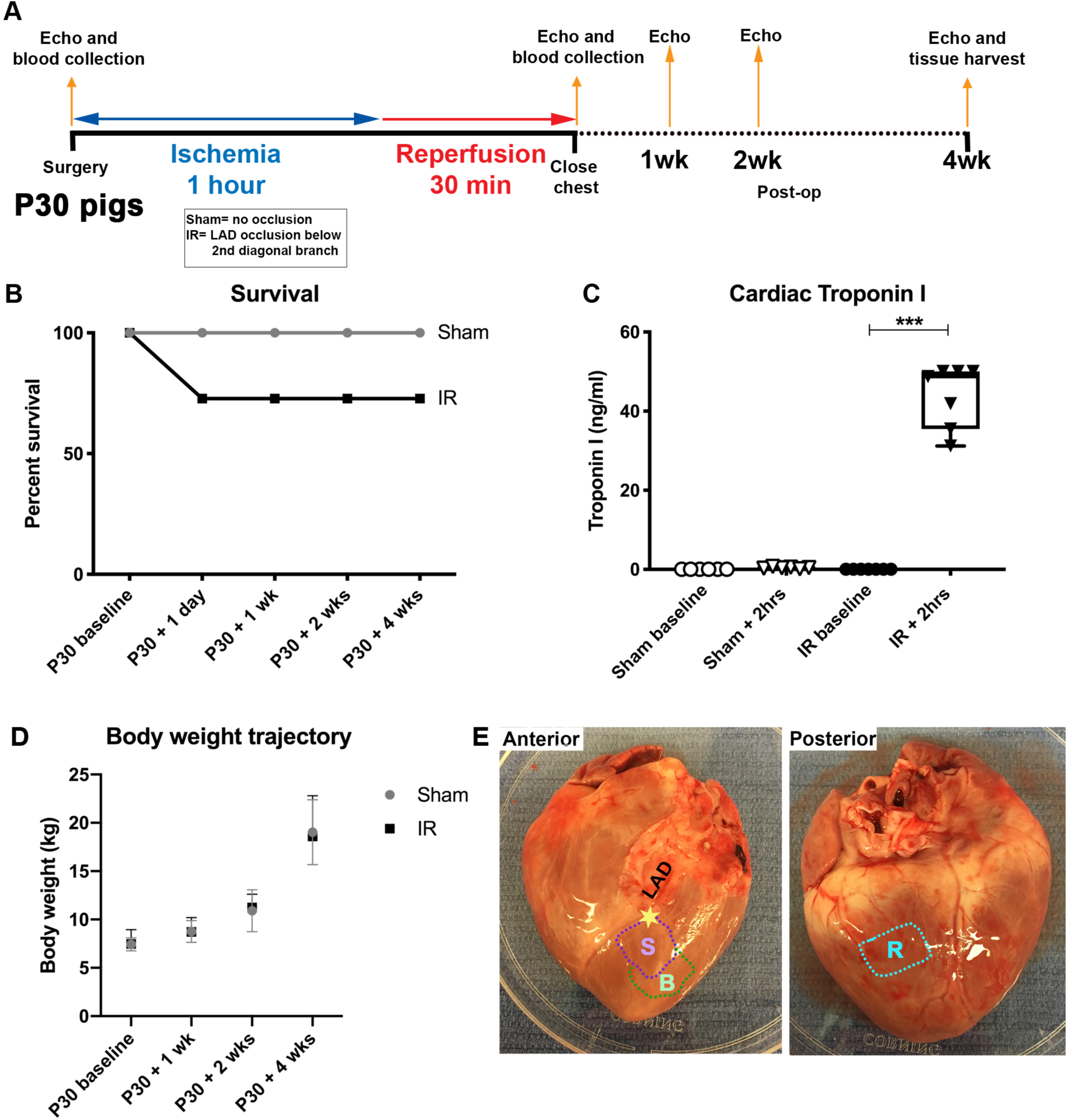
Cardiac ischemia/reperfusion injury assessment in P30 pigs. Schematic representation (A) of P30 pig ischemia/reperfusion (IR) procedure. Ischemia was induced for 1 hour by occlusion of the left anterior descending artery (LAD) just distal to the second diagonal branch. Pigs were monitored for 30 minutes prior to closing the chest. Weekly/biweekly echocardiography (echo) assessments were undertaken up until tissue harvest at 4 weeks post-surgery. (B) Survival shown as a percent of the whole cohort in sham and IR pigs was assessed at regular intervals over the course of the study. (C) Cardiac Troponin I (ng/ml) was measured from blood plasma at baseline and 2 hours following the initiation of ischemia or sham surgeries, to assess the extent of cardiac injury. (D) Body weights (kg) were measured prior to surgery, and then at regular time points post-surgery to assess growth trajectories of sham-operated and IR pigs over the 4 week study. (E) Image of a representative heart at 4 week tissue harvest, showing anticipated scar zone (S) and border zone (B) in relation to the occlusion site (represented as a star) just below the second diagonal branch of the LAD, and remote zone (R) taken from the posterior left ventricle. Data are mean ± SD. 2-way ANOVA with Tukey post-hoc analysis (B&D) and Kruskal-Wallis test with Dunn’s post-hoc analysis (C) were performed as appropriate, ***p<0.001, n=6-7/group Troponin I, n=8/group body weight.

**Table 1:** Cohorts of pigs used in the cardiac injury procedure. The number of P30 pigs used in both sham-operated and ischemia/reperfusion (IR) study groups, alongside interventions carried out during surgery and key observations made at tissue harvest 4 weeks post-surgery.

|  |  | Sham | IR |
| --- | --- | --- | --- |
| **Total number operated** |  | 8 | 11 |
| **Survival (number) at 4 weeks post-surgery** |  | 8 | 8 |
| **Number of females** |  | 4 | 4 |
| **Number of males** |  | 4 | 4 |
| **Intervention during surgery** | **Epinephrine administration** | None | n=5 (2 successful) |
|  | **Defibrillation** | None | n=5 (2 successful) |
| **Number with adhesions present around heart at harvest** |  | 2 | 7 |
| **Body weight (kg)** |  | 19.03±3.35 (NS) # | 18.60±4.21 (NS) |
| **Heart weight (g)** |  | 136.10±25.06 (NS) | 166.20±46.08 (NS) |

Serum cTnI was increased >40-fold 2 hours following ischemic injury in IR pigs compared with the sham-operated animals (Figure 1C), demonstrating an effective myocardial injury. Sodium levels declined and potassium levels increased, in both sham and IR pigs at 2 hours post-surgery, with lactate and glucose levels increasing only in IR pigs, consistent with ischemic injury (Supplementary Table 2). Together these blood indicators, along with ST-segment changes on ECG, and observed myocardial color change, confirm myocardial tissue damage and ischemic injury in pigs subjected to the IR surgery.

No differences in body weight growth trajectories for the two groups was found over the course of the study protocol (Figure 1D). In addition, heart weight and body weight measurements at tissue harvest were similar between groups (Table 1). Thus, the ischemic cardiac injury, via temporary LAD occlusion, did not affect overall growth or development of the animals. At the time of sacrifice when collecting the tissue during harvest, it was noted that a greater number of pigs within the IR group had substantially denser pericardial adhesions compared with the sham group (Table 1). The hearts were excised and left ventricular myocardium was isolated with scar, border and remote zones collected as shown in Figure 1E. The scar zone was defined as immediately distal to the LAD suture site in both sham and IR pigs, and thus contained injured and viable myocardium. This IR protocol in P30 pigs produces a limited myocardial injury without significant morbidity or mortality of the operated animals.

### Cardiac function declines post-ischemic injury in P30 pigs

Echocardiographic assessment of cardiac function showed a decline in ejection fraction (EF, Figure 2A) and fractional shortening (FS, Figure 2B) in IR pigs from pre-operative baseline to 2 hours post-injury. EF was similarly decreased at 1 and 2 weeks in IR compared to sham-operated animals. By the study endpoint, at 4 weeks post-surgery, EF and FS were unchanged compared with the assessment 2 hours post-surgery for IR or sham-operated groups. Thus, the decrease in cardiac function was apparent by 2 hours following IR, but did not improve or decrease further by 4 weeks post-injury.

**Figure 2:**
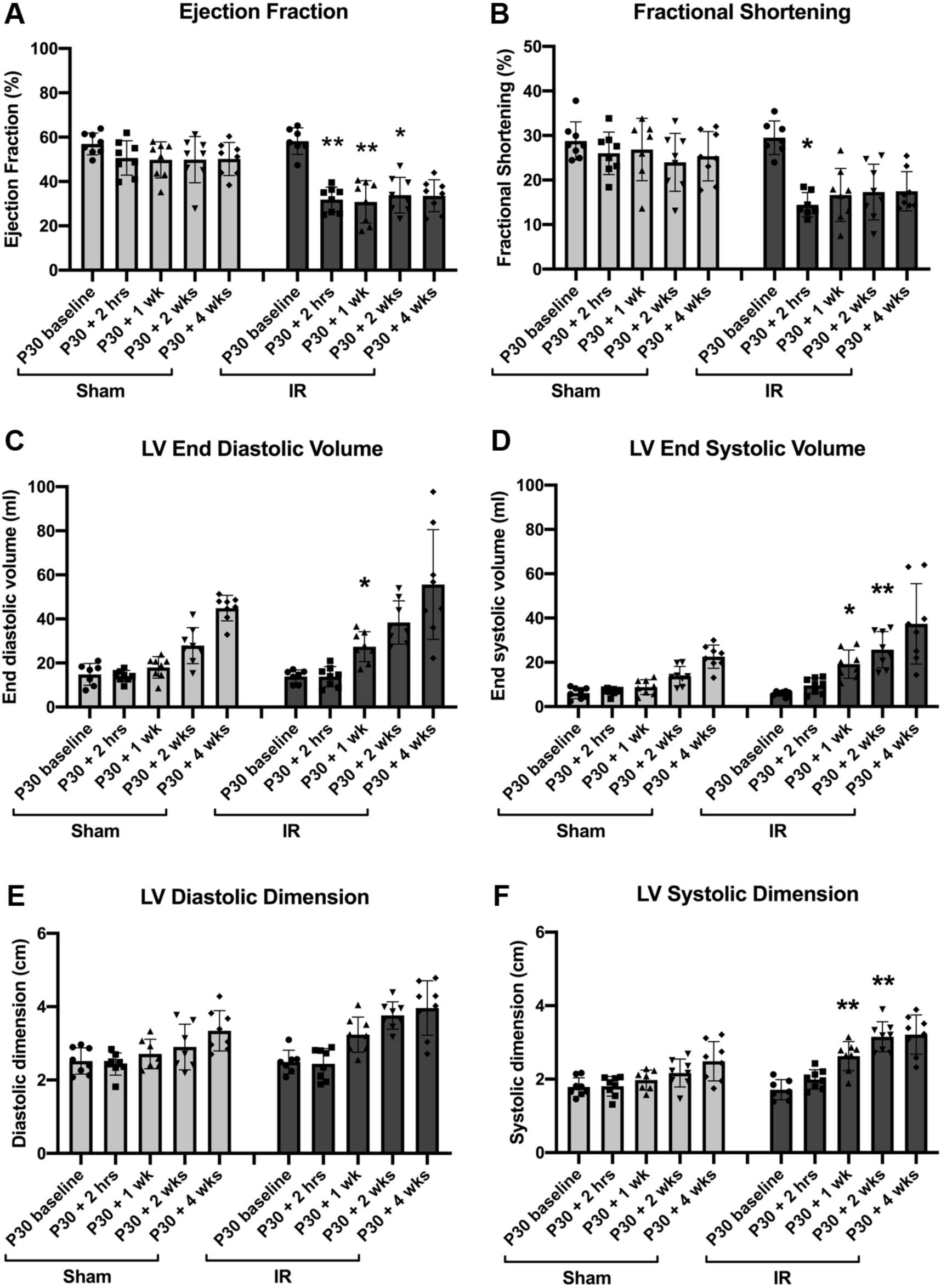
Cardiac function decreases at 2 hours post-ischemia/reperfusion (IR) injury, but does not decline up to 2-months of age. Echocardiography analysis of sham-operated and IR pigs measured prior to surgery (P30 baseline), 2 hours following the start of ischemia (or sham equivalent), 1 week, 2 weeks, and 4 weeks following surgery. (A) Ejection Fraction (%), (B) Fractional Shortening (%), (C) Left Ventricular (LV) End Diastolic Volume (ml), (D) LV End Systolic Volume (ml), (E) LV Diastolic Dimension (cm) and (F) LV Systolic Dimension (cm). Data are mean ± SD. Mixed effect ANOVA analysis with Bonferroni’s post-hoc test *p<0.05, **p<0.01 vs same time point in sham, n= 8/group.

Corresponding changes were observed in LV volumes and dimensions in systole and diastole (Figure 2C-F). Likewise, ventricular septum and LV posterior wall thicknesses were unchanged between sham and IR pigs (Supplementary Table 3). Assessment of LV size by echocardiography showed the expected growth increase as the pigs age. Stroke volume initially dropped post-injury, however was then similar between sham and IR pigs throughout the study, with heart rate being within the normal range (Supplementary Table 3). These results indicate that, while cardiac function worsens following IR in P30 pigs, heart growth parameters are in the normal range.

### Cardiomyocyte hypertrophy is not induced with IR injury, in the month after surgery

Cardiac hypertrophy after IR injury was assessed by measurement of organ size and CM area. Heart weight corrected for body weight, 4 weeks post-surgery, in IR pigs was increased compared with sham-operated or unoperated pigs (Figure 3A). However, there were no significant differences in heart weights or body weights when analysed separately between sham-operated and IR pigs (Table 1). In addition, histological assessment of CMs showed that cross sectional area of individual CMs was unaltered between unoperated control, sham and IR pigs, as well as amongst three different zones of the left ventricular myocardium (Figure 3B, C). Thus, CMs were of a similar size in all 2-month-old pigs heart assessed, with and without ischemic injury. This is further confirmed with no changes in mRNA expression of hypertrophic genes *NPPA* and *NPPB* between unoperated, sham and IR pig hearts (Figure 3D). No evidence of hypertrophic growth following injury was also corroborated with no change in the posterior wall thickness of the LV between sham and IR pigs, via echocardiography assessment (Supplementary Table 3).

**Figure 3:**
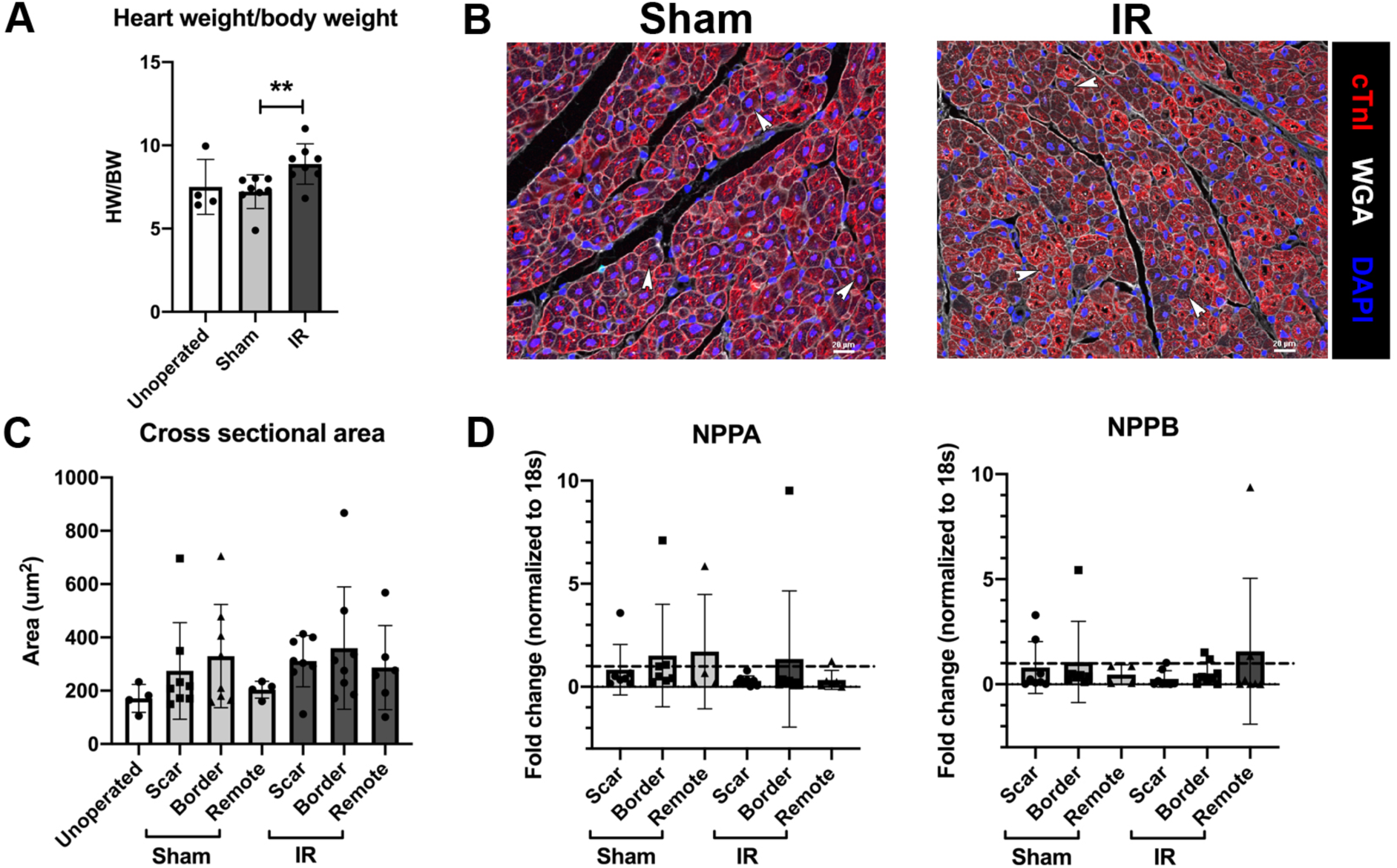
Cardiomyocytes (CMs) do not show hypertrophic growth 4 weeks after ischemia/reperfusion (IR) injury. (A) Heart weight corrected for body weight assessment in unoperated 2-month-old controls, sham-operated and IR pigs at tissue harvest. (B) Representative myocardial scar region sections stained with cardiac Troponin I (cTnI, red), DAPI (blue) and wheat germ agglutinin (WGA, white), used for assessing cardiomyocyte cross sectional area (CSA), with arrowheads indicating examples of CM with central nuclei used for counting. (C) Quantification of CM CSA 4 weeks following surgery in unoperated controls, sham-operated and IR pigs divided into zonal regions: scar, border and remote. RT-qPCR for hypertrophic genes (D) *NPPA* and *NPPB* mRNA in sham and IR pigs, with separated zones, normalized to *18S* and shown relative to unoperated 2-month control myocardial mRNA (dashed line). Data are mean ± SD. Mann-Whitney test (A) **p<0.01, 2-way ANOVA with Tukey post-hoc analysis (C&D; NS), n=4-8/group. Scale bar=20μm.

Gene expression levels of fetal and adult sarcomeric isoform genes were also investigated. Expression of the fast myosin heavy chain 6 (*MYH6*), characteristic of immature cardiac sarcomere in pigs (22, 52), was increased in IR pig myocardium, particularly within the scar region versus sham pigs (Supplementary Figure 2A). However, the immature slow skeletal type troponin I1 (*TNNI1*, encoding ssTNI), as well as *TNNI3 and* slow *MYH7*, characteristic of mature pig CMs, showed no change between sham and IR pigs (Supplementary Figure 2B-D). Upon immunohistochemical staining for the immature ssTNI, similar expression was observed in the myocardium in both sham and IR pigs, across all three zones. However, expression was most obvious at the epicardial region in each zone, with a gradual decrease towards the endocardium (Supplementary Figure 2E). Our results therefore show that following injury there is an increased expression of *MYH6*, suggestive of a partial reactivation of fetal sarcomeric gene expression, compared with sham-operated pigs, but other sarcomeric proteins are largely unaffected.

### Cardiomyocyte cell cycling activity is unchanged following P30 ischemia/reperfusion cardiac injury

In unoperated pigs at 2-months-of-age, mitotic activity as identified by phospho-histone H3 (pHH3)-positive nuclei, is apparent in CMs as they become multinucleated (Supplemental Figure 1A) (50). Similar cell cycling in multinucleated CMs was observed in sham-operated or IR hearts, as measured by immunohistochemical staining of pHH3 with CMs identified by cardiac troponin I (Figure 4A). Interestingly, multinucleated CMs with each individual nucleus stained positive for pHH3 were found in sham, IR, and unoperated control cohorts (Figure 4A and Supplementary Figure 1A). To determine CM mitotic index, pHH3-positive CMs were assessed in cross-section, to count one pHH3 positive nucleus per cell (Figure 4B and Supplementary Figure 1B). No difference between sham and IR pigs was shown for CM cell cycling activity (Figure 4C), or between myocardial zones, suggesting that IR cardiac injury does not cause a change in CM proliferation. Thus, CM cell cycling is active in 2-month-old pig hearts, as indicated by pHH3 reactivity in CM nuclei, however cell cycle activity was unchanged between sham-operated and IR pigs.

**Figure 4:**
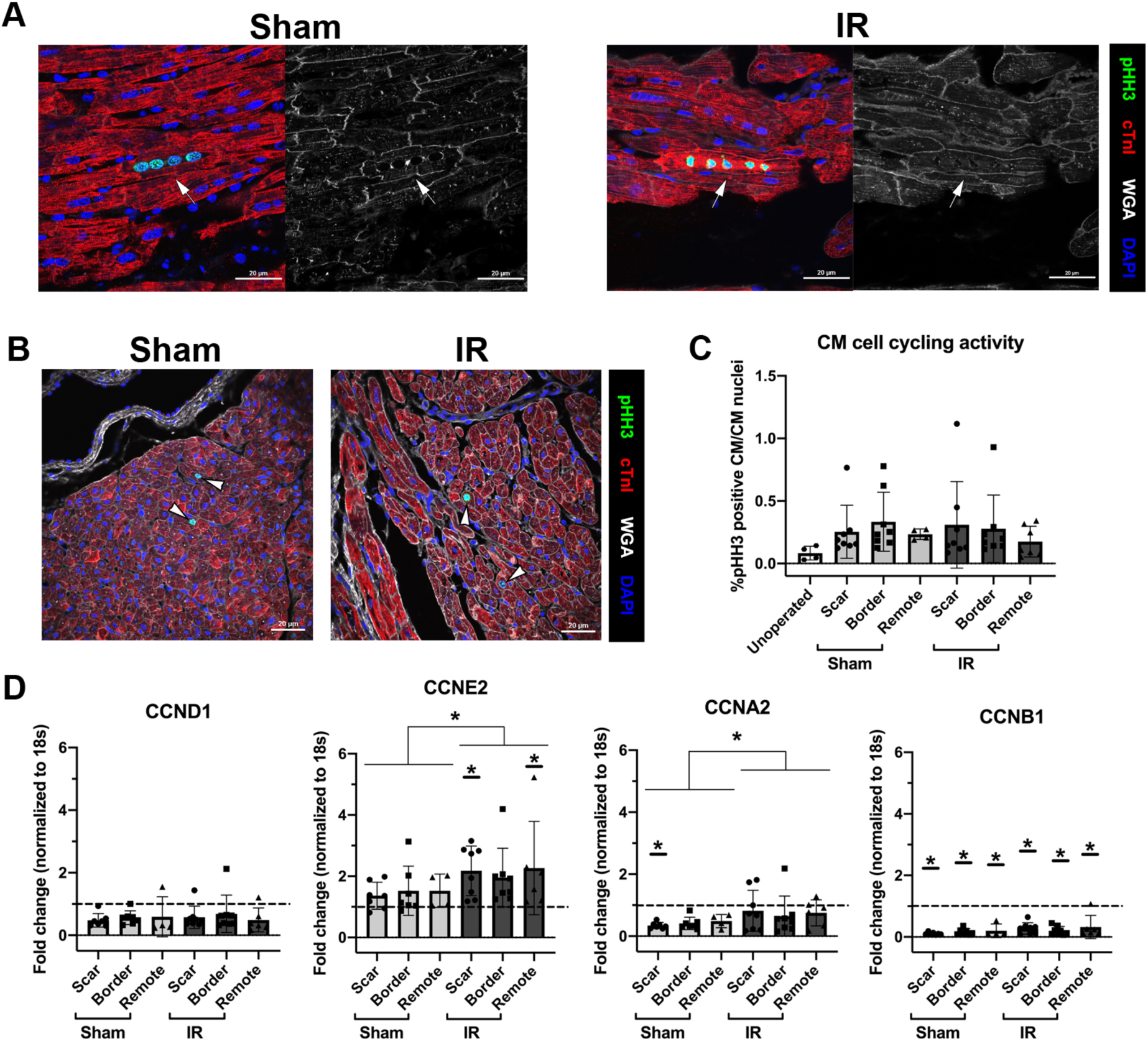
Cardiomyocytes (CMs) are multinucleated with cell cycling similar between sham and IR pigs at 4 weeks post cardiac surgery. (A) Immunohistochemistry of sham and IR scar regions of left ventricular myocardium at 2-months-of-age shown in the longitudinal orientation, highlighting multinucleation, as depicted by cardiac Troponin I (cTnI, red) in combination with DAPI (blue) with wheat germ agglutinin (WGA, white). In individual CMs with cell cycling activity multiple nuclei show pHH3 immunoreactivity (green), indicated by the arrow. (B) Representative cross-section images of CM cell cycling in sham and IR (scar zones) pigs used for counting are shown with pHH3-positive CM nuclei indicated by white arrowheads. (C) Quantitation of %pHH3-positive CM, relative to total CM nuclei identified by cTnI+DAPI in unoperated control, sham and IR pigs (3 zones). (D) RT-qPCR mRNA expression of cell cycling genes *CCND1*, *CCNE2, CCNA2* and *CCNB1* in sham-operated and IR pigs, with separated zones (scar, border, remote), normalized to *18S* and shown relative to unoperated 2-month control myocardial mRNA (dashed line). Data are mean ± SD, with C&E analyzed by 2-way ANOVA with Tukey post-hoc analysis (C; NS. E *p<0.05). Underlined * indicates significant difference vs unoperated controls (*p<0.05). n=4-8/group. Scale bar=20μm.

Expression of cell cycle-associated genes was examined by RT-qPCR of mRNA from myocardial tissue zones of sham-operated and IR pigs. mRNA expression of the cell cycling genes *CCNE2* (present during G1/S phase of the cell cycle) and *CCNA2* (activated during S phase) was increased in all zones together in IR pigs compared with sham (Figure 4D). *CCNE2* was also increased versus unoperated 2-month-old control pig hearts (Figure 4D). However, *CCND1* (activated during G1), *CCNA2* and *CCNB1* (activated during M phase of the cell cycle) showed a decline in expression of both sham and IR hearts versus unoperated 2-month-old controls (Figure 4D). This indicates that following cardiac IR injury, there may be slightly increased cell cycling activity and that sham and IR may have differences in cell cycling relative to unoperated controls. However, coordinated increases in multiple cyclins needed for active progression through the cell cycle were not observed. As whole tissue was taken for mRNA assessment, it cannot be confirmed whether or not any changes in cell cycling activity occurs in CMs or other cell types (e.g. fibroblasts or immune cells). It is important to note that these hearts were collected 4 weeks following surgery so any immediate effect of injury on CM proliferative capacity would not be detected if transient. However, taken together, mRNA assessment of cell cycling 4 weeks post-surgery suggests there may be minor changes in DNA replication following injury, though these do not equate to an overall change in CM proliferative activity.

### Vessel density and cell death are unchanged following P30 ischemia/reperfusion cardiac injury

Vascular density and cell death were assessed in the myocardium 4 weeks following injury as indicators of tissue damage and cardiac repair. Microvessel density, determined in lectin-DAB stained sections of the myocardium, was similar between unoperated control, sham-operated, and IR pigs at 2-months-of-age (Supplementary Figure 3A, B and Supplementary Figure 1D). When the epicardial region was specifically assessed, there appeared to be a thickening of the epicardium with a lack of vascular density in both sham and IR pigs (Supplementary Figure 3C). A high level of variability was seen within IR pigs in epicardial vascular density (Supplementary Figure 3D), likely due to variable injury between individual pigs. Cell death, as assessed by the apoptotic marker TUNEL, was measured to determine whether or not cardiac injury had a damaging effect on cell survival. Cell death was unchanged within the CM and non-CM region of the myocardium between unoperated control, sham-operated, and IR pigs (Supplementary Figure 1C and 3E, F). Thus, IR in P30 pigs did not alter vascularity or myocardial cell death when assessed 4 weeks following injury.

### The immune response is upregulated in IR pigs, particularly within the scar region

To determine the differential gene expression changes between sham-operated and IR pigs, myocardial RNA was analyzed by RNA-sequencing (RNA-seq). Multiple comparisons were undertaken including overall sham versus IR, followed by each group split based on scar or border zones and also based on sex (Supplementary Figure 4A). Of particular interest were the gene expression changes within the expected scar region in sham-operated versus IR pigs. GO term analysis revealed that biological processes typically related to an immune response were upregulated within the IR scar zone versus a comparable area in sham pig myocardium (Figure 5A). Such upregulated terms consist of immune system processing, response, and regulation (including *CD69, CD72, CD86, CD209, IL10, IL18, IL18R1*), as well as cell surface receptor signaling pathways and response to stimuli (including *CXCL9, CXCL10, CXCL11, CXCR3*). These main GO terms of differential gene expression associated with immune response were restricted to the ‘scar’ zone, as relatively fewer differences were found within the border zone of sham versus IR pigs (Supplementary Figure 4A). Thus, the majority of RNA transcriptional changes associated with inflammation in IR pigs occur close to the site of injury.

**Figure 5:**
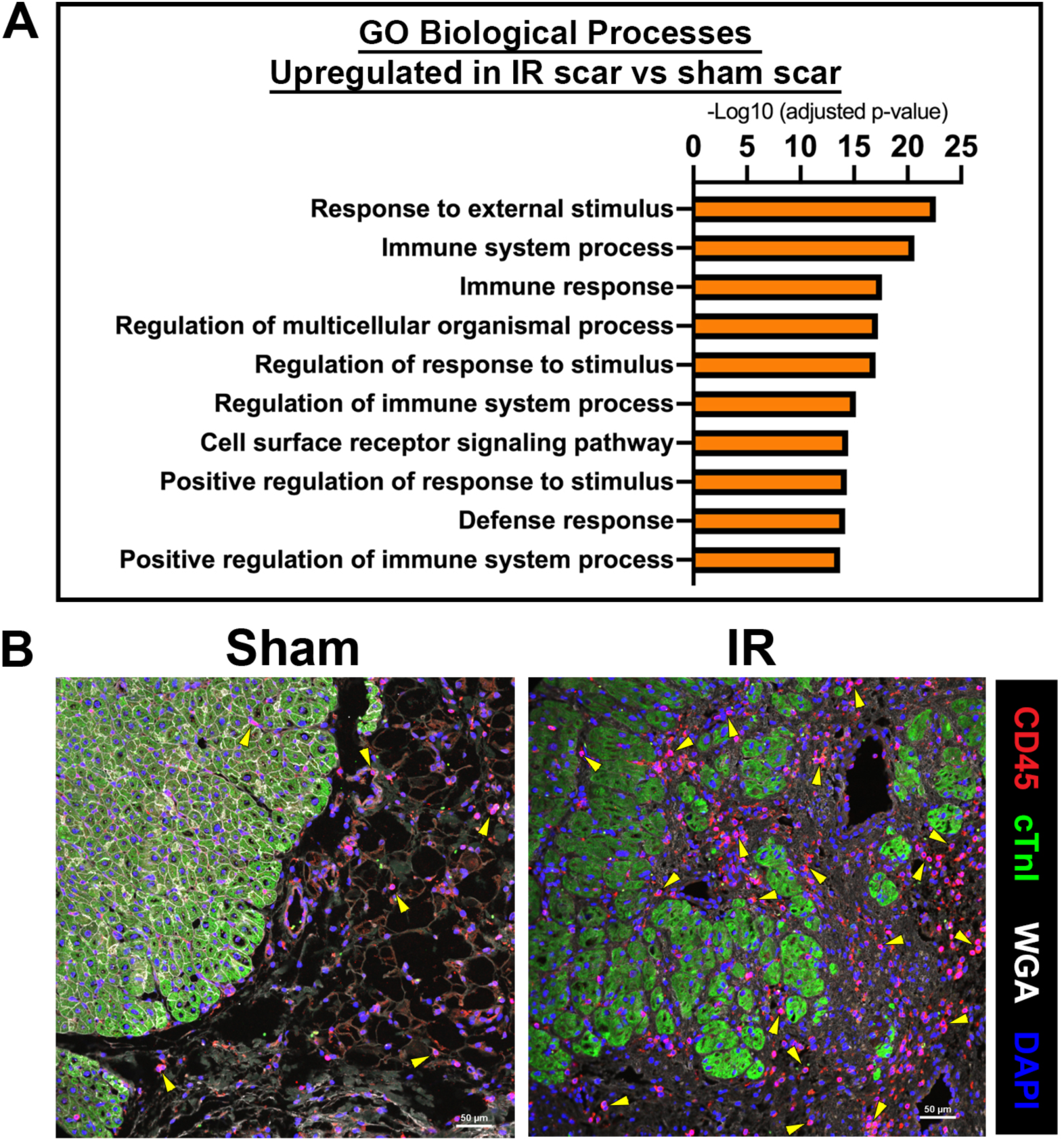
The immune response is upregulated in the scar region of IR pigs 4 weeks following ischemic cardiac injury. RNA-sequencing was undertaken on total RNA from sham and IR scar zones. (A) Gene ontology enrichments for biological processes that are upregulated in IR versus sham-operated pigs within the scar zone highlights an immune response to injury, n=4 per group. (B) Representative images of sham-operated and IR scar regions (at the epicardial region with immunofluorescence for CD45-positive immune cells (red) alongside myocardium (cTnI, green), wheat germ agglutinin (WGA, white) and nuclei (DAPI, blue). Yellow arrowheads represent examples of CD45-positive immune cells. Scale bar=50μm.

Immunohistochemical staining for CD45+ immune cells confirmed an upregulation of the immune response within the scar zone in IR pigs compared with sham pigs (Figure 5B). Following histological analysis, more CD45+ immune cells were visualized within the epicardial region of the scar in IR pigs compared to sham pigs. CD45 stains for all leucocytes, and there were a substantial number evident in the epicardial region, as well as in scar zone, of IR pig hearts 4 weeks following cardiac ischemic injury. Overall, RNA-seq and histological analysis shows increased immune response and presence of CD45+ cells 4 weeks following IR cardiac injury.

### Myocardial scarring is apparent 4 weeks following IR cardiac injury in P30 pigs

To determine whether cardiac ischemic injury at P30 resulted in scar formation, myocardial samples were assessed for gene expression levels of extracellular matrix (ECM) related genes as well as histological staining. From RNA-seq, GO term analysis revealed that the cellular components associated with ECM regulation were upregulated in IR scar versus sham scar zone tissue (Figure 6A). However, the RNA-seq findings of ECM upregulation in IR scar tissue were not validated when assessing mRNA expression levels of individual ECM genes in an increased number of animals. No differences in expression between sham and IR cohorts or myocardial zones were found for ECM related genes (Supplementary Figure 4B-I). Genes typically expressed during scar formation, *COMP* and *CHAD*, showed variable expression levels in both sham and IR pigs (Supplementary Figure 4H&I). These findings highlight that sham-operated pigs with thoracotomy and exposure of the LAD showed a response to the cardiac surgery, even without LAD occlusion, as mRNA expression levels in both sham and IR were altered versus unoperated controls. Differences between the RNA-seq dataset and mRNA expression levels was likely due to smaller sample sizes being assessed via RNA-seq compared with all pigs for mRNA, leading to increased variability. Thus, significant changes in ECM mRNA expression levels were not detected by RT-qPCR between sham-operated and IR animals.

**Figure 6:**
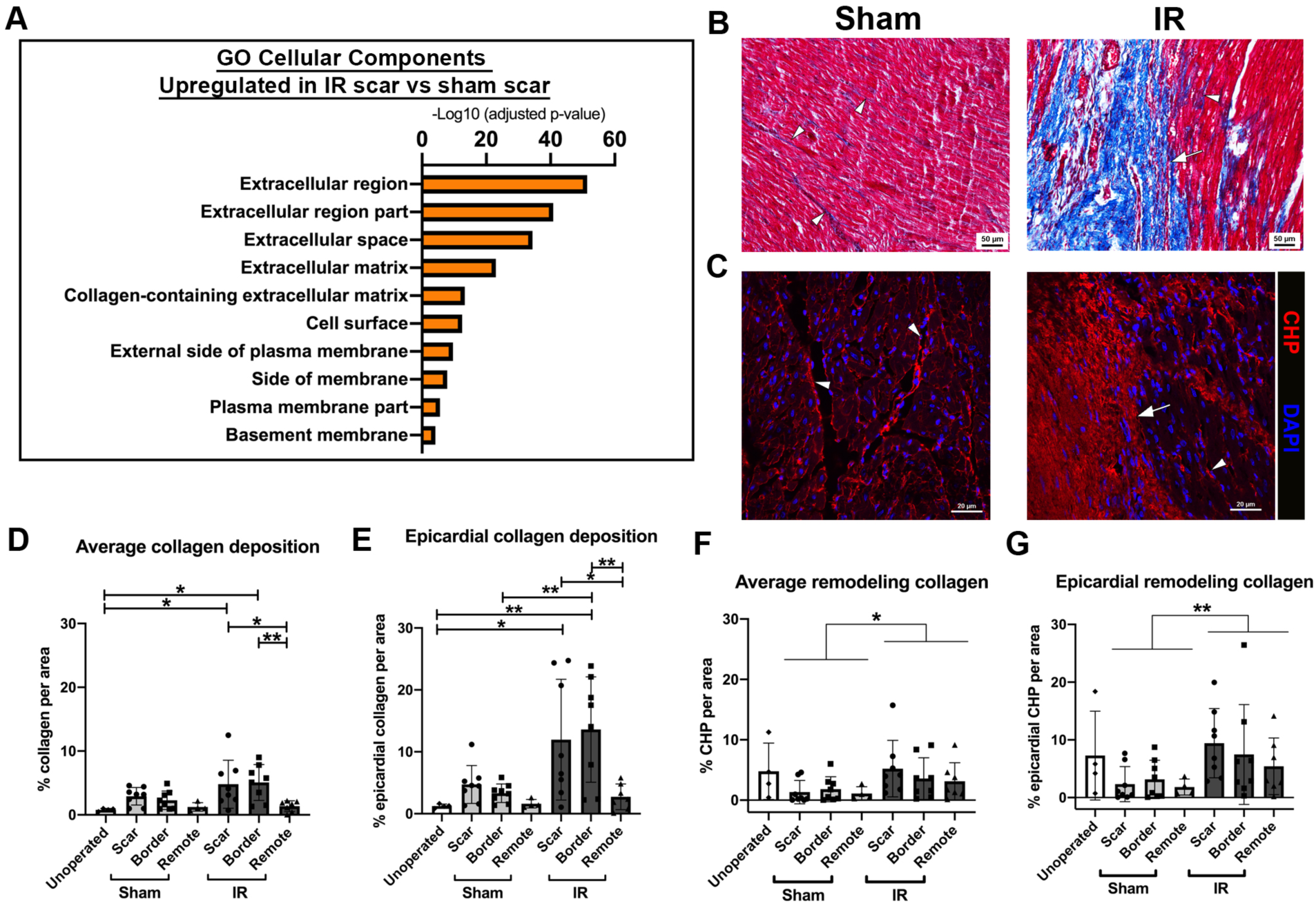
Scar formation is evident at 4 weeks post-surgery in ischemia/reperfusion (IR) pigs. RNA-sequencing was undertaken on total RNA from sham and IR scar zones. (A) Gene ontology enrichments for cellular components that are upregulated in IR versus sham-operated pigs, within the scar zone, showing ECM gene expression changes in response to injury. (B) Representative Masson’s Trichrome staining (myocardium=pink, collagen=blue), in sham-operated and IR scar region sections. The arrow depicts obvious scar formation in IR pigs, with arrowheads showing interstitial collagen in sham-operated and IR myocardial sections. (C) Representative images in sham and IR scar zone of immunohistochemistry for Collagen Hybridizing Peptide (CHP, Helix3), marking remodeling collagen (red) alongside DAPI for nuclei (blue). The arrow depicts dense scar area, with arrowheads indicating interstitial remodeling collagen (red). (D) Quantification of average collagen deposition as indicated by Masson’s trichrome staining at 2-months-of-age across all zones of sham and IR pigs, as well as unoperated control myocardial tissue sections. Collagen (blue staining) was measured relative to tissue area (μm^2^) and displayed as a percentage. (E) The epicardial region of each zone was also quantified (similar to D), as this was the area of most prominent scar formation proximal to the LAD ligation. (F) Quantification of average CHP deposition at 2-months-of-age across all 3 zones of sham and IR pigs, as well as unoperated control myocardial tissue sections. CHP (red) was measured relative to tissue area (μm^2^) and displayed as a percentage. (G) The area of CHP reactivity per total area in the epicardial region of each zone was also quantified. Data are mean ± SD. 2-way ANOVA analysis with Tukey post-hoc analysis, *p<0.05, **p<0.01 n=4-8/group. Scale bar=50μm (B), 20μm (C).

Scar formation and collagen remodeling were assessed after IR injury. Histological analysis of fibrillar collagen by Masson’s trichrome staining scar zones in all pigs demonstrated an obvious scar at the epicardial region in IR pigs, that was not observed in sham-operated animals (Figure 6B). When multiple images per zone of the myocardium were stained for collagen deposition, collagen accumulation was increased by >5-fold in the scar and border zone of IR pig hearts, compared to unoperated 2-month old control pigs (Figure 6D; Supplementary Fig 1E). As expected, the remote zone of IR pigs showed less collagen deposition than the scar and border zones. The most prominent scarring was obvious at the epicardial region of the myocardium, thus, when this area was quantified, there was an increase in collagen deposition in IR pigs versus sham (Figure 6E). As expected, the remote zone had less collagen deposition than the epicardial region, consistent with greater myocardial injury in the anterior region of the heart proximal to the LAD occlusion. Although not significant, trending increases in collagen deposition in sham-operated animals (Figure 6D, E) may be indicative of mild cardiac injury during the process of opening the chest cavity and placing a suture in the myocardium (without occlusion). However, large regions of collagen deposition at the site of injury were only observed in animals subjected to IR.

Immunohistochemical staining for Collagen Hybridizing Peptide (Helix3), for detection of remodeling collagen, illustrated a similar obvious scar formation in IR pigs versus sham (Figure 6C, F). Again, the greatest change in ECM response between sham and IR pigs, as indicated by remodeling collagen, was apparent in the epicardial region of the myocardium (Figure 6G). 2-month-old unoperated control pigs showed a variable level of remodeling collagen, hence no significant differences between this cohort and the surgical groups were observed (Supplementary Figure 1F). Taken together, IR injury via LAD occlusion in P30 pigs creates obvious scar generation with ECM remodeling at 4 weeks post-IR. This is most apparent at the area closest to the occlusion site predominant on the epicardial surface.

### Sex differences in response to cardiac surgery are observed in young pigs

Female and male pigs were used in this study to identify any sex differences in the response to IR cardiac injury. As these pigs were P30 at time of surgery, they were considered to be pre-pubertal (39). By echocardiography, cardiac functional parameters, EF and FS revealed sex differences, however only within the sham cohort (Figure 7A, B). Female sham-operated pigs overall had a lower EF and FS than males after surgery. Thus, females responded differently to males with a more substantial drop in cardiac function following the chest being opened and a suture being placed around the LAD (without IR).

**Figure 7:**
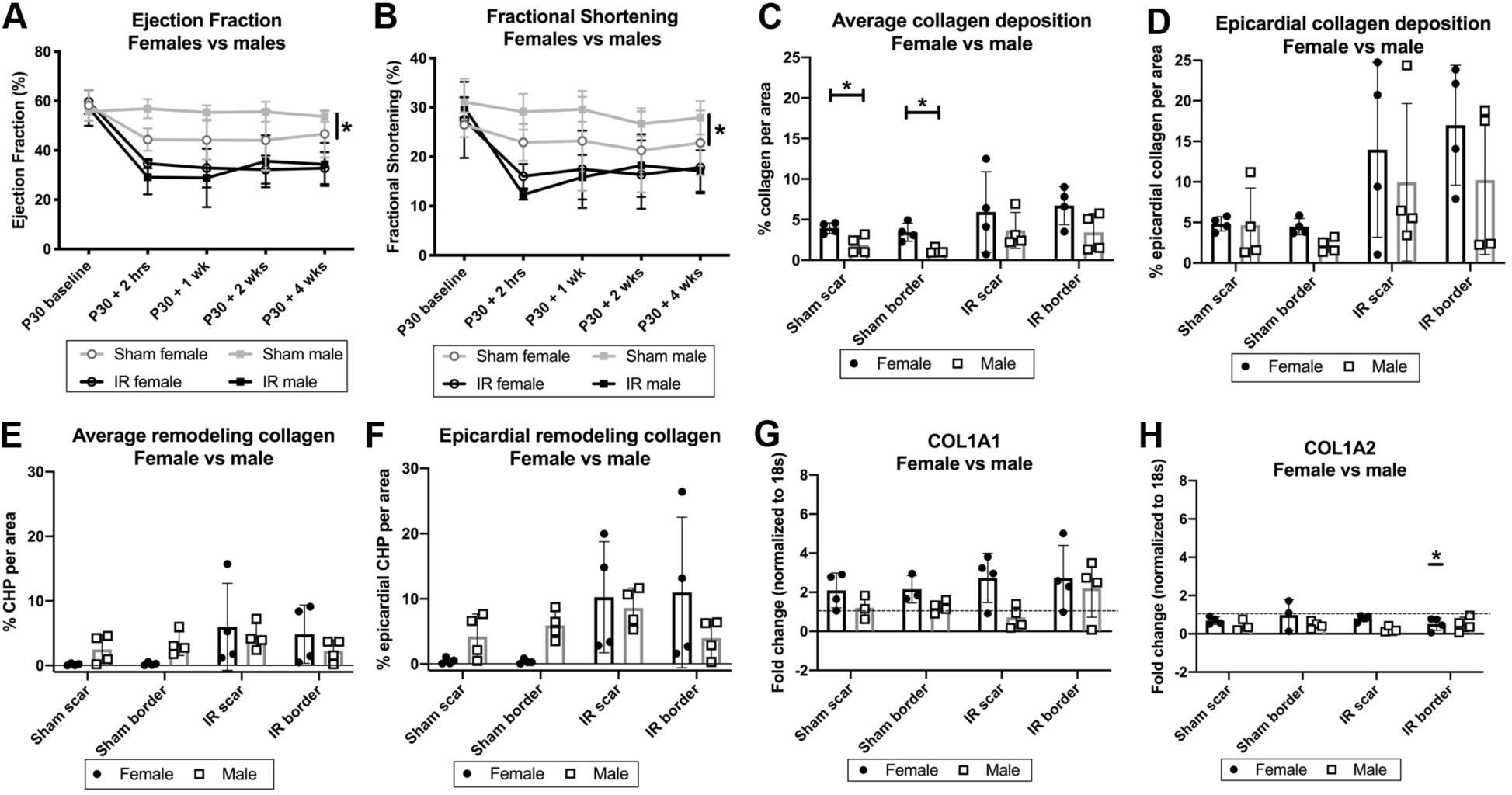
Sex differences are observed, particularly within the sham-operated cohort, when assessing ECM changes. Cardiac functional parameters (A) Ejection Fraction (%) and (B) Fractional Shortening (%) were calculated separately for females and males in each surgical group at each timepoint by echocardiography from baseline to 4 weeks post-surgery. The areas of Masson’s trichrome staining indicative of total collagen (C, D) and Collagen Hybridizing Peptide (CHP) immunohistochemical staining (E, F) were also calculated separately for females and males. (C) Female and male sham-operated and IR scar and border zones for quantification of average collagen deposition, shown as collagen deposition relative to tissue area (μm^2^) and displayed as a percentage. (D) The epicardial region of each group as shown in C, was also split to asses sex differences in the area with greatest scar formation. (E&F) CHP quantification also split based on sex, similar to C&D. RT-qPCR of ECM-related mRNA changes between sexes for (G) *COL1A1* and (H) *COL1A2* in female and male sham-operated and IR pigs, with separated zones (scar and border), normalized to *18S* and shown relative to unoperated 2-month control myocardial mRNA (dashed line). Underlined * = significant from unoperated controls. Data are mean ± SD, 2-way ANOVA with Tukey post-hoc analysis for C-H and Mixed effect ANOVA analysis with Bonferroni’s post-hoc for A&B. *p<0.05, n=3-4/group.

Sex differences related to ECM changes were detected histologically 4 weeks after injury, with female pigs showing a greater response to surgery via increased collagen deposition within the scar and border zones of the heart, as compared to males (Figure 7C). This was also confirmed via mRNA expression changes in *COL1A1* and *COL1A2* with higher levels in female pigs (Figure 7G, H). This higher fibrotic response in females was particularly shown within the sham-operated cohort. However, when remodeling collagen was assessed, opposite sex differences were shown. Male pigs showed a greater CHP immunoreactivity in the sham-operated cohort versus females (Figure 7E, F). In contrast, sham-operated females had greater overall collagen deposition as indicated by Masson’s trichrome staining, particularly in sham pigs, suggesting that male pigs may still be undergoing ECM remodeling. However, the response to ischemic injury is similar between sexes, with possible trends for females to create a larger response. Overall, our results suggest female pigs undergoing surgery may have a worse outcome with lowered cardiac function and greater fibrotic response than males.

## DISCUSSION

While there is extensive information available on postnatal heart development and regenerative potential after injury in rodents (12, 28, 34, 35, 43), much less is known about cardiac repair and regeneration in large mammals, including humans (49). Here, we report cardiac repair following ischemia/reperfusion injury in P30 pigs, with both sexes represented. We demonstrate that P30 pigs lack the capacity of cardiac regeneration following IR, despite appreciable mitotic activity in multinucleated CMs in normal pigs at this stage (50). Myocardial damage is evident immediately following IR injury, with a decrease in cardiac function detected 2 hours after injury, that does not improve or worsen over the 4 week study period. While cell cycling in multinucleated CMs is evident in unoperated, sham-operated and IR pigs at 2-months-of-age, CM cell cycle activity is not increased at 1 month post-injury. Cardiac repair is evident in the pigs subjected to IR with the formation of a localized scar and increased inflammation in the epicardial region proximal to the region of ischemic injury. Interestingly, indications of sex differences in cardiac function and scarring also were observed. This study therefore demonstrates that transient ischemic injury in the hearts of young swine leads to decreased cardiac function and increased scarring, but not increased proliferation of CMs.

The temporary occlusion of the LAD, immediately distal to the second diagonal branch, results in a significant decrease in cardiac function, as measured by ejection fraction and fractional shortening 2 hours after surgery. Such a decrease in function is similar to previously reported swine models of LAD ligation and permanent myocardial infarction (MI) (5, 6, 36, 44, 45, 48, 51, 54). The decreased cardiac function after IR was then maintained to 4 weeks post-injury, with no obvious morbidity or mortality beyond the day of the surgery. In an adult pig model of IR, ejection fraction is maintained at a comparable decreased level from baseline to 2 months post-surgery (44). By 3 months post-IR there is a further decrease in cardiac function, alongside CM hypertrophy, suggesting heart failure progression. In contrast, IR injury in P30 pigs did not lead to changes in chamber wall morphology or CM cross sectional area, suggesting a lack of pathologic cardiac hypertrophy 4 weeks following surgery. Future studies are required to determine if these young pigs develop heart failure 3 months post-IR, as has been reported in adult swine (44). This transient cardiac injury model therefore is effective in showing that even a mild injury (of 1 hour ischemia) can lead to decreased cardiac function, which does not worsen over time, in addition to a cardiac healing response apparent in inflammation and scarring.

When assessed at 2-months-of-age, unoperated, sham-operated and IR pigs in this study showed evidence of CM mitotic activity (pHH3 expression) and multinucleation. This contrasts with a previous report that pig CMs exit the cell cycle before P30 (54), however confirms findings from our lab that show CM cell cycling and multinucleation are active at 2-months-of-age (50). In mice, CMs undergo binucleation, cell cycle arrest, and lose their capacity to regenerate approximately 1 week after birth. In contrast, CM cell cycling and mononucleation persist at 2 months in pigs. However, severe cardiac injury by permanent MI after P3 in pigs does not produce a regenerative response (51, 54). Moreover, here we show that a milder transient cardiac ischemic injury at P30 also does not lead to increased CM cell cycling 4 weeks following surgery, even though CM nuclei undergoing multinucleation exhibit mitotic activity at the time of injury. Whether this injury approach alters CM cell cycling activity in the initial stages following IR is yet to be determined. Nevertheless, CM mitotic activity in young swine is not sufficient to produce cardiac regeneration beyond a few days after birth, similar to what has been reported for rodents.

The cardiac injury response in young swine after the postnatal period includes inflammation and scarring. Young pigs after cardiac injury produce an immune response to initiate cardiac healing and subsequent ECM remodeling, as has been observed in older large and small mammals (30, 37, 51, 54). With transient ischemic injury in P30 pigs, we observed localized ECM upregulation with increased inflammation and thickening of the epicardial collagen matrix at the site of injury. Similar epicardial activation and thickening has been observed in adult mice and humans after cardiac ischemic injury and may be part of the mammalian cardiac injury response (7, 53). Here we show that a mild transient ischemic injury at P30 in pigs also leads to localized scarring at the site of injury. In contrast to permanent MI after P3, the scar generated in pigs at P30 did not span the entire myocardium, however, the transient IR injury created a small regional scar in hearts with reduced EF observed within 2 hours post-IR. Since reduced cardiac function was observed within hours after injury, the damage to the myocardium itself that did not recover after injury is likely to be the major contributor to reduced cardiac output. In addition, scar generation and ECM remodeling apparent after cardiac injury may be inhibiting any reparative/regenerative response to cardiac injury beyond the postnatal period in mammals.

There is increasing attention to potential sex differences in cardiovascular biology and disease in studies of animal models and human patients (17, 23, 31, 38). In the current study, we evaluated the effects of cardiac IR injury on female and male pigs prior to sexual maturity (39). While IR-specific differences in cardiac function were not observed between females and males, the female pigs subjected to surgery and thoracotomy in sham-operated group exhibited reduced cardiac function and increased collagen expression relative to males. These data provide initial evidence that pre-pubertal female and male pigs may respond differently to cardiac surgery, with pre-pubertal female pigs being predisposed to a greater detrimental response. In many studies of cardiac injury response in mice and humans, males are predominantly studied due to their greater fibrotic response to cardiac injury (16, 17, 27). In adult females of reproductive age, estrogen signaling is antifibrotic and cardioprotective after cardiac injury (14, 15, 33), but may be less effective prior to puberty or after menopause. As the pigs in this study are pre-pubertal, it is likely that females do not yet have high enough circulating estrogen levels to create a protective effect after cardiac injury, leading to adverse outcomes compared with males. Since sex-differences are observed in studies of cardiac injury response, it is important that both female and male animals are included in large mammalian cardiac preclinical injury studies.

The ability of the human infant heart to regenerate after injury is not known, but there is an anecdotal report of a newborn infant suffering MI with full functional recovery after thrombolytic therapy (13). Another report showed that infants with anomalous origin of the left coronary artery from the pulmonary artery who had surgical repair at 10 months-of-age, had significant cardiac functional recovery with little/no evidence of scar formation, as assessed by cardiac magnetic resonance imaging (20). This suggests that humans, like mice and pigs, have a postnatal capacity for cardiac regenerative repair, but the duration of cardiac regenerative potential in humans is not known (8, 13, 26, 29). While pigs are comparable in size to humans with a similar neonatal developmental trajectory, cardiac physiology, and coronary vessel architecture (24, 41), CM nucleation, ploidy and growth characteristics are not conserved among mammalian species. In addition, there is increasing evidence that other factors may contribute to loss of cardiac regenerative capacity in mammals soon after birth. Our results, together with previous reports, show that cardiac injury after the postnatal period of P0-P3 leads to inflammation and scarring, not CM renewal in young pigs, despite presence of cell cycling CMs. Loss of cardiac regenerative capacity has been linked to oxidative stress at birth, ECM maturation, and infiltration of macrophages in addition to CM cell cycle exit (11, 19, 34, 51, 54). Further understanding the factors which inhibit regeneration in the developing pig will add to the current limited knowledge on mechanistic regulation of cardiac repair following injury in young large mammals. This improved knowledge, through use of models as described in this study, will aid in improving and testing mechanisms to reactivate and promote heart regeneration in adults.

## ACKNOWLEDGMENTS

We thank members of the Yutzey lab for valuable input and discussion. We also are grateful for the support from Veterinary Services at CCHMC, especially Angie Cummins. We also thank Dr. Inigo Valiente Alandi for his assistance with RNA-seq data analysis, and Dr Steve Houser and his team for pig protocol advice. We thank Chrissy Schulte from the animal imaging laboratory for her assistance with echocardiography.

