## Supplemental Methods, tables and figures for "SCAR FORMATION AND DECREASED CARDIAC FUNCTION FOLLOWING ISCHEMIA/REPERFUSION INJURY IN 1-MONTH-OLD SWINE"

#### Immunohistochemistry

Sections were dewaxed with xylene (Fisher Scientific, Hampton, NH) and rehydrated in decreasing concentrations of ethanol (Decon Labs Inc, PA). Antigen retrieval was carried out in Tris-EDTA buffer (pH 9; Abcam) using a pressure cooker. Slides were blocked with 6% goat serum, followed by staining with Troponin I Type 1 (slow skeletal) Antibody (ssTnI, 1:200, NBP1-56641, Novus Biologicals) at 4°C overnight. Fluorophore-conjugated secondary antibody (1:400; Alexa Fluor ab175471, Abcam, Cambridge, UK) was used to detect immunostaining, with the addition of Wheat Germ Agglutinin (WGA), Alexa Fluor™ 647 Conjugate (1:250; Thermo Fisher Scientific, Waltham, MA) and DAPI (5mg/ml, D1306; Thermo Fisher Scientific). Stains were imaged using a Nikon Eclipse Ti Fluorescence microscope with NIS elements software (Tokyo, Japan).

#### Lectin-DAB Staining

Glycoprotein lectin was used to identify capillary endothelial cells in all myocardial sections. Sections were dewaxed, rehydrated and antigen retrieval carried out as above. Slides were blocked with 0.3% $H_2O_2$, followed by 6% goat serum. Sections were stained with Biotinylated Lectin Antibody (B-1105, Vector Laboratories, Burlingame, CA) in a 1:300 dilution. Following overnight incubation, mouse Avidin-Biotin Complex (ABC; Thermo Scientific) was added to the stained sections. 3,3'-Diaminobenzidine (DAB) was the chromogen used for visualization of the lectin signals, made using DAB metal concentrate (10X) in stable peroxide substrate buffer (1X) at a 1:100 dilution (Thermo Scientific). Stains were imaged using an Olympus BX51 microscope and Nikon DS-Ri1 camera with NIS elements software. Microvessel density, measured as small vessel counts per tissue area ($\mu m^2$), was calculated using an automated analysis program created on NIS Analysis elements software.

#### Cell death detection

Cell death was detected by TUNEL using the *In Situ* Cell Death Detection Kit, Fluorescein (Roche, Basel, Switzerland). Briefly, slides were baked for 1 hour following by dewaxing and rehydration. Antigen retrieval was achieved via proteinase K addition (20μg/ml; Carlsbad, CA). Slides were blocked in 4% goat serum, then incubated overnight with the monoclonal Anti-α-Actinin (Sarcomeric) antibody (A7811, 1:100)(Millipore Sigma), before the addition of TUNEL enzyme (1:10, Roche), Goat Anti-Mouse IgG H&L (Alexa Fluor® 568) (1:400, ab175473, Abcam) and Wheat Germ Agglutinin, Alexa Fluor™ 647 Conjugate (1:250; Thermo Fisher Scientific). Cell death was imaged as above, and quantified as the number of cardiomyocytes (identified via sarcomeric α-actinin stain) and non-cardiomyocytes normalized to total nuclear number (DAPI).

### TABLES

**Supplementary Table 1: Porcine primer sequences used for RT-qPCR analysis of pig myocardial mRNA normalized to *18S* ribosomal RNA.**

| **RNA** | **Sequence 5'-3'** | |
|---|---|---|
| | **Forward** | **Reverse** |
| *18S* | AATTCCGATAACGAACGAGACT | GGACATCTAAGGGCATCACAG |
| *NPPA* | TGAACCCAGCCCAGAGAGAT | CAGTCCACTCTGTGCTCCAA |
| *NPPB* | GTTGCTGCTAGGATGCCGTT | TACCTCCTGAGCACATTGCAGC |
| *CCND1* | GCGAGGAACAGAAGTGCG | TGGAGTTGTCGGTGTAGATGC |
| *CCNE2* | TCAAGACGCAGTAGCCGTTT | AGCCAAACATCCTGTGAGCA |
| *CCNA2* | CCCTGCATTTGGCTGTGAAC | ATTCAGGCCAGCTTTGTCCC |
| *CCNB1* | CATGCAGGATAATTGTGTGCCC | CCTCGATTCACCACGACGAT |
| *MYH6* | GTGAAGAGATAACCAGAGGAGCG | CACCTGATCCTCCTTCACGG |
| *MYH7* | AAGGTCAAGGCCTACAAGCG | CTTTGTTGCGCCCTCAGGAT |
| *TNNI1* | AACTTCACGCCAAGGTGGAG | ATGGCCTCGACGTTCTTTCT |
| *TNNI3* | ATACGACGTGGAGGCGAAAG | CATCATGGCATCGGCAGAGA |
| *COL1A1* | CTGGAAGAGCGGAGAATACTG | CTGTAGGTGAAGCGGCTGTT |
| *COL1A2* | CTTCGTGCCTAGCAACATGC | CAAAGTTCCCGCCAAGACCA |
| *COL3A1* | CCTGGACGAGATGGAAACCC | GGCTACCTACTGCACCTTGG |
| *FN1* | ACCCTTGCAGTTCCGAGTTC | TCCCTGACGATCCCACTTCT |
| *POSTN* | GTGATCCACGGAGAGCCAATTA | ATGACCATCGCCACCTTCAAT |
| *TNC* | TCGCTACAAGCTGAAGGTGG | ACCAGTTGACACCCTGACTG |
| *COMP* | CGACTACGCGGGTTTCATCT | GAACCGCACTCTGATGTAGC |
| *CHAD* | ACGCTGAAACACGTCCATCT | TCAGGACCTGTTTAATGGCGA |

**Supplementary Table 2: iSTAT blood analyzer parameters at baseline versus 2 hours post-surgery for sham and ischemia/reperfusion (IR) surgery groups.**

| | **P30 baseline** | | **P30 + 2 hours** | | ***% change between time points*** | |
|---|---|---|---|---|---|---|
| **Parameter** | **Sham #** | **IR** | **Sham** | **IR** | ***Sham*** | ***IR*** |
| Sodium | 134.50±1.73 | 137.00±1.41 | 131.50±2.52** | 132.50±2.38*** | *-2.23% *** | *-3.28% **** |
| Potassium | 4.13±0.45 | 4.48±0.73 | 5.68±0.17** | 6.03±1.28** | *+37.53% *** | *+34.60% *** |
| Calcium | 1.51±0.07 | 1.48±0.11 | 1.48±0.03 | 1.47±0.14 | *-1.99%* | *-0.68%* |
| Lactate | 2.00±0.51 | 1.60±0.56 | 2.30±0.53 | 2.78±0.91* | *+15%* | *+73.75% ** |
| Glucose | 102.50±9.54 | 81.00±6.88 | 125.25±29.88 | 138.50±9.33** | *+22.20%* | *+71.00% *** |

**#** Data are mean ± SD and were analyzed by Mixed effect ANOVA analysis with Bonferroni's post-hoc tests. P values indicate significant differences compared to same surgical group at baseline (highlighted in bold), *p<0.05, **p<0.01, ***p<0.001, n=4 for all groups, with calcium: n=4 sham, n=5 IR and lactate: n=3 sham, n-4 IR. No significant differences were observed for sham vs IR cohorts.

**Supplementary Table 3: Tissue Doppler Echocardiography measurements of P30 pigs subjected to sham or ischemia/reperfusion (IR) surgery, measured at baseline (prior to surgery), 2 hours, 1 week, 2 weeks, and 4 weeks after surgery in 2D four-chamber and M-mode images.**

| | | | | | | |
|---|---|---|---|---|---|---|
| **Parameter** | **Surgery** | **P30 baseline** | **P30 + 2 hours** | **P30 + 1 week** | **P30 + 2 weeks** | **P30 + 4 weeks** |
| **Ventricular Septum Diastolic Thickness (cm)** | **Sham #** | 0.43±0.04 | 0.41±0.09 | 0.51±0.12 | 0.59±0.07 | 0.59±0.12 |
| | **IR** | 0.48±0.10 | 0.43±0.06 | 0.47±0.20 | 0.55±0.07 | 0.58±0.14 |
| **Ventricular Septum Systolic Thickness (cm)** | **Sham** | 0.70±0.15 | 0.61±0.15 | 0.76±0.10 | 0.93±0.14 | 0.90±0.27 |
| | **IR** | 0.79±0.14 | 0.53±0.06 | 0.72±0.24 | 0.70±0.15 | 0.76±0.26 |
| **LV Posterior Wall Diastolic Thickness (cm)** | **Sham** | 0.44±0.06 | 0.47±0.03 | 0.46±0.08 | 0.56±0.09 | 0.56±0.09 |
| | **IR** | 0.42±0.04 | 0.44±0.06 | 0.47±0.11 | 0.55±0.14 | 0.55±0.17 |
| **LV Posterior Wall Systolic Thickness (cm)** | **Sham** | 0.62±0.10 | 0.66±0.07 | 0.74±0.08 | 0.80±0.10 | 0.88±0.13 |
| | **IR** | 0.67±0.10 | 0.68±0.11 | 0.75±0.18 | 0.81±0.21 | 0.88±0.20 |
| **Stroke Volume (ml)** | **Sham** | 8.27±2.86 | 7.14±1.70 | 9.21±2.44 | 14.09±5.70 | 22.35±3.38 |
| | **IR** | 8.16±2.23 | 4.27±1.38 ** | 8.23±2.69 | 12.66±3.14 | 18.24±8.09 |
| **Heart Rate (bpm)** | **Sham** | 145.28±20.46 | 107.22±10.05 | 177.50±20.84 | 147.44±27.85 | 131.11±10.59 |
| | **IR** | 149.72±28.44 | 126.52±34.15 | 183.71±27.90 | 171.14±13.37 * | 147.62±18.11 |

**#** Data are shown as mean ± SD, with Mixed-Effect ANOVA Model with post-hoc Bonferroni tests * $p<0.05$, ** $p<0.01$ vs sham surgery at equivalent time. All parameters shown are significantly different over time as the pigs age (via mixed effect model analysis for repeated measures, thickness parameters $p<0.01$, stroke volume and heart rate $p<0.0001$). n=4-8/group.

### FIGURES

#### Supplementary Figure 1

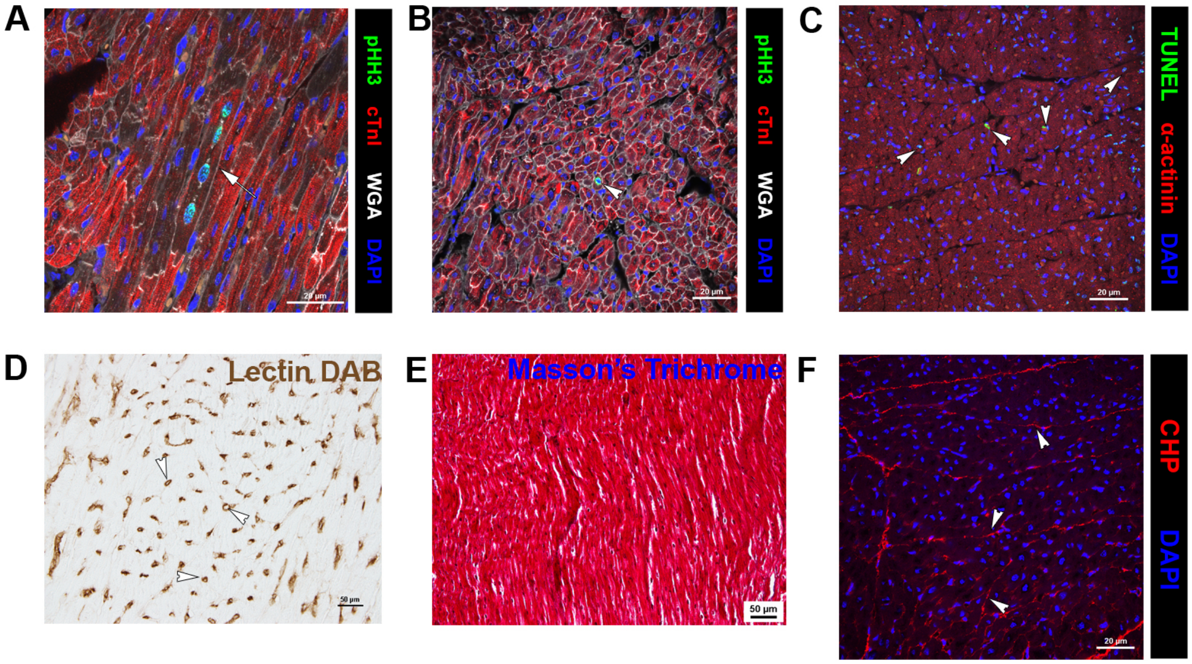

**Supplementary Figure 1: Control unoperated 2-month-old pig heart histology, shows evidence of cardiomyocyte (CM) multinucleation, cell cycling, and cell death, with minimal interstitial collagen deposition.** Representative histological images for 2-month-old unoperated control pig hearts, showing (A) multinucleated CM in long-axis with mitotic activity [pHH3 (green) immunohistochemistry in cardiac troponin I (cTnI, red) stained CM, with wheat germ agglutinin (WGA, white) staining plasma membranes and DAPI (blue) nuclei]. (B) cross sectional representation of (A) used for pHH3 positive CM counting. Example images of (C) cell death in sarcomeric $\alpha$-actinin (red) stained CM as indicated by TUNEL staining (green), (D) microvessel density, via lectin-DAB staining, (E) interstitial collagen deposition (blue), via Masson's trichrome stain (myocardium pink), and (F) interstitial remodeling collagen [as measured by Collagen Hybridizing Peptide (CHP, red)]. Arrowheads represent example cells used for counting or areas of interstitial collagen quantified. Images shown are representative of n=4. Scale bar=20$\mu$m (A-C, F), 50$\mu$m (D,E).

### Supplementary Figure 2

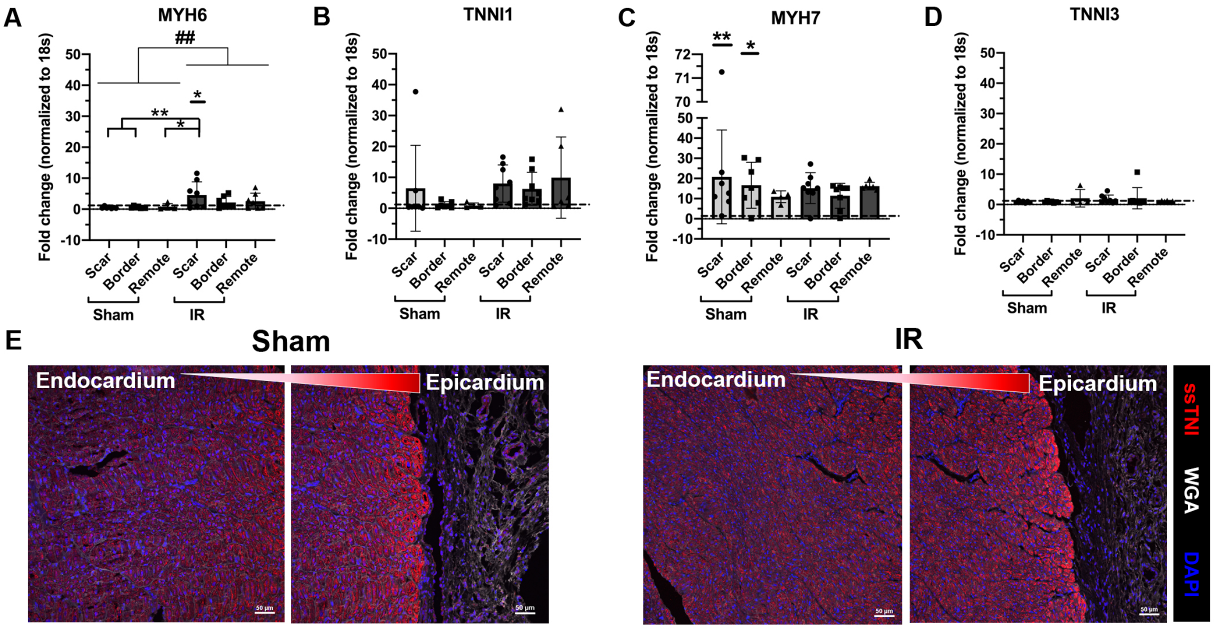

**Supplementary Figure 2: Following ischemia/reperfusion (IR) cardiac injury, higher expression of *MYH6* mRNA is observed in the myocardium compared to sham and unoperated 2-month-old control pigs.** Expression levels of fetal and adult sarcomeric protein isoform genes were evaluated by RT-qPCR normalized to *18S* and shown relative to unoperated 2-month control myocardial mRNA (dashed line). Fetal genes (A) *MYH6*, and (B) *TNNI1,* compared to adult isoforms (C) *MYH7,* and (D) *TNNI3,* are shown in sham and IR scar, border and remote regions from total RNA collected 4 weeks following surgery. (E) Representative immunofluorescent staining of ssTnI (encoded by *TNNI1*) with wheat germ agglutinin (WGA, white) staining plasma membranes and DAPI (blue) staining nuclei, with the arrow indicating visually higher levels of ssTnI at the epicardium diminishing nearer the endocardium. (A-D) Data are mean $\pm$ SD, 2-way ANOVA (## $p<0.01$) with Tukey post-hoc analysis, *$p<0.05$, **$p<0.01$, underlined * = significance compared to unoperated controls. n=4-8/group. Scale bar=50μm.

**Supplementary Figure 3**

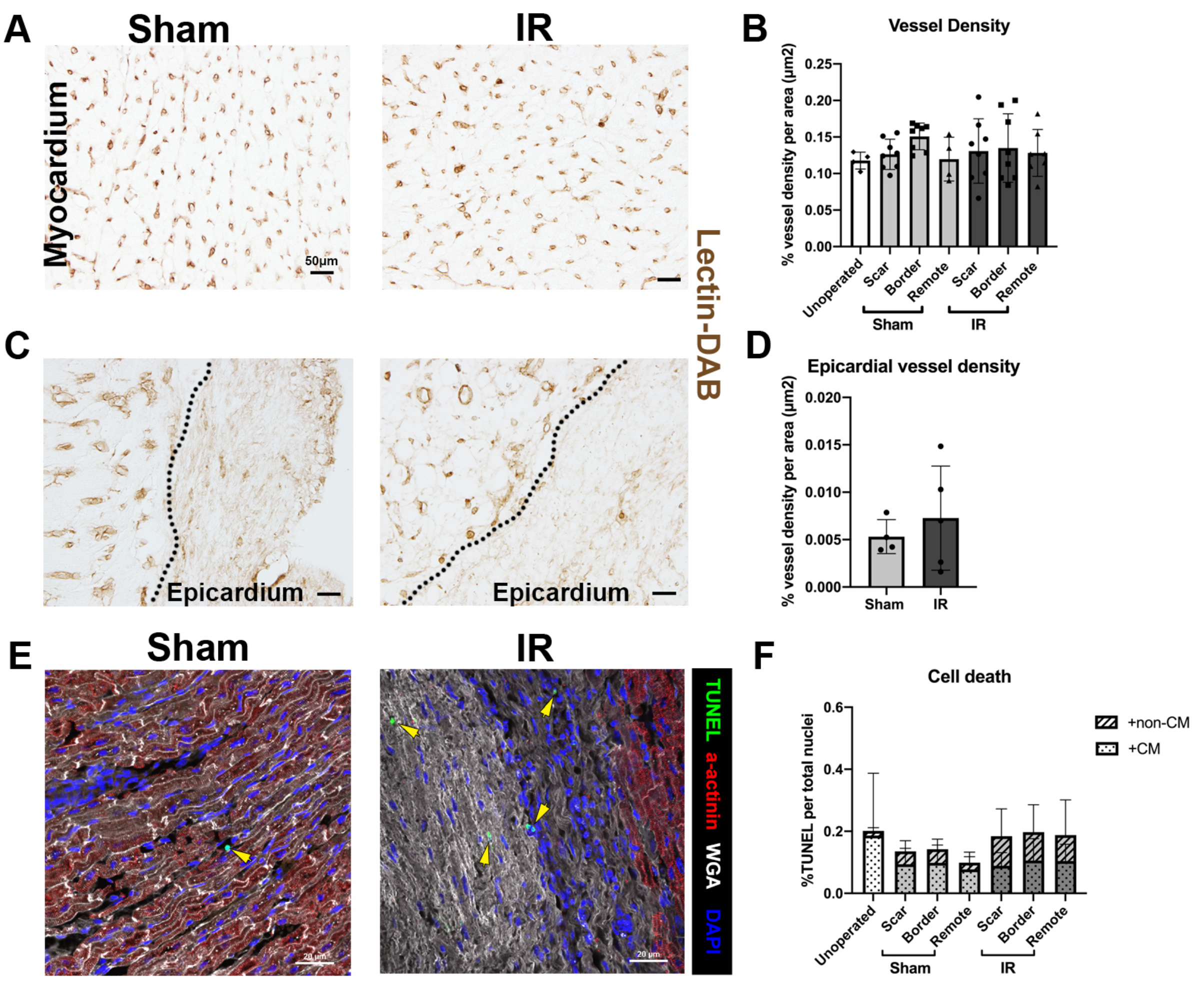

**Supplementary Figure 3: Vascular density and cell death are similar between sham and IR pigs at 4 weeks post-cardiac surgery.** (A) Representative images of lectin-DAB stained microvessels in sham and IR myocardium (scar zone), with (B) quantification of unoperated control, sham, and IR (scar, border, and remote zones) pig vascular density, represented as % vessel density (lectin-DAB staining) per tissue area ($\mu m^2$). (C) epicardial regions of the hearts were stained for lectin-DAB to assess vascular changes within the region of most scar formation and (D) quantified in sham and IR pig scar zones. Dotted line represents edge of myocardium. (E) Representative images of cell death, as indicated by immunohistochemical TUNEL staining (green) with sarcomeric $\alpha$-actinin (red) stained CM (yellow arrow heads, non-CM TUNEL positive cells), in sham and IR pigs (scar zones). (F) Quantitation of cell death in unoperated, sham, and IR (scar, border, and remote zones), expressed as a percentage of TUNEL positive cells per total nuclei (DAPI) and split based on the CM area (nuclei within cTnI positive region) and non-CM area (nuclei within cTnI negative region). Data are mean ± SD. 2-way ANOVA with Tukey post-hoc analysis (B&F) or Mann-Whitney tests (D) were performed as appropriate with no significant differences detected. n=4-8/group. Scale bar=50$\mu$m (A,C), 20$\mu$m (E).

### Supplementary Figure 4

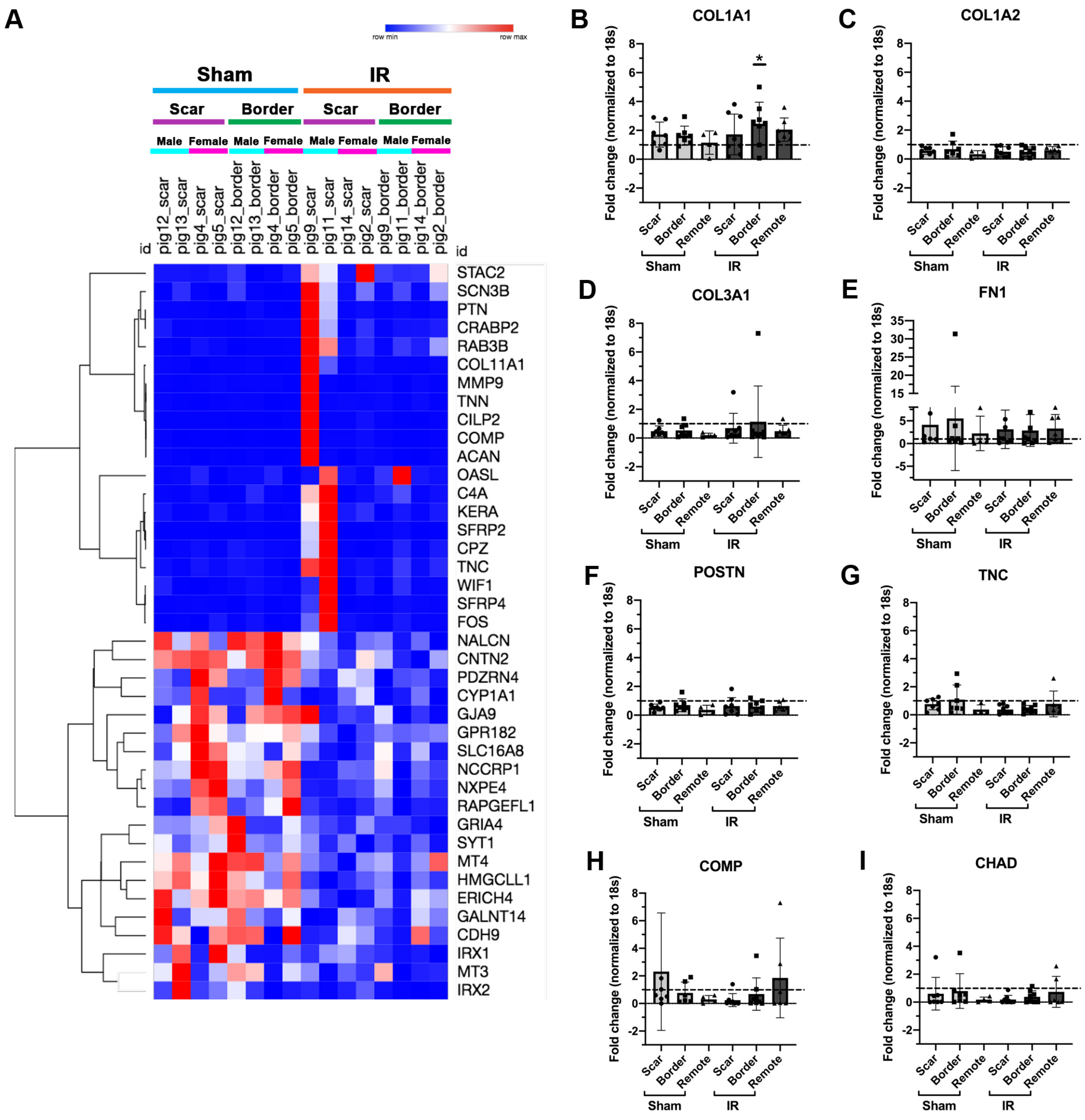

**Supplementary Figure 4: RNA-seq analysis suggests gene expression changes between sham and IR pigs, but significant differences were not observed by RT-qPCR analysis of mRNA expression of ECM related genes.** (A) Heatmap of top 20 up and down-regulated genes from sham and IR pigs (n=8), scar and border zones (n=4) for female and male pigs (n=2). Expression levels of individual ECM genes were determined by RT-qPCR normalized to *18S* ribosomal RNA and shown relative to unoperated 2-month control (n=4) myocardial mRNA levels (dashed line). Sample sizes are n=7-8 for all sham and IR groups, apart from sham remote n=4. ECM genes analyzed are (B) *COL1A1* (C) *COL1A2*, (D) *COL3A1,* (E) *FN1* (F) *POSTN*, (G) *TNC,* (H) *COMP* and (I) *CHAD.* Data are mean ± SD, 2-way ANOVA with Tukey post-hoc analysis, *$p<0.05$, underlined * = significant from unoperated controls.
